# Computational Platform for streamlining the success of sequential antibiotic therapy

**DOI:** 10.1101/2025.02.16.638553

**Authors:** Alejandro Anderson, Matt Kinahan, Rodolfo Blanco-Rodríguez, Alejandro H. Gonzalez, Klas Udekwu, Esteban A. Hernandez-Vargas

## Abstract

The scarcity of antibiotics and the need for swift decision-making present significant challenges for healthcare practitioners. When confronted with such circumstances, practitioners must prioritize their approach based on several key factors. By leveraging the recent biological discovery of collateral sensitivity, we have devised an open-source computational platform. This platform utilizes the drug resistance profile of an evolved strain and initial population conditions to potentially predict the failure of cycling therapy in eradicating or restraining a multi-drug resistant bacterial population. We demonstrate how this framework can anticipate potential failures for a range of antibiotics in chronic *pseudomonas aeruginosa* infections. This innovative methodology lays the foundation for evolutionary therapies that can facilitate the selection of appropriate treatments, thereby reducing antibiotic resistance.

## INTRODUCTION

The silent pandemic of antibiotic resistance is a pressing global health crisis causing a high death toll ***Levy (1998***); ***Mayers et al. (2017***); ***Levy and Marshall (2004***); ***Murray et al. (2022***). The discovery of new classes of antibiotics is urgently needed. However, the development of antibiotics may not keep up with the escalation of antibiotic resistance in the next few decades ***Gold and Moellering Jr (1996***); ***Annunziato (2019***); ***Bollenbach (2015***).

Based on this scenario, a foundational interrogative would be if we can develop strategies for better treatment by using our current arsenal of antibiotics. Addressing this question would require the effective forecasting of bacterial evolution to personalized therapies known as “Evolutionary Therapies” ***Perry (2021***). Examples of this could be scheduling the order of antibiotics to take advantage of predictable patterns of bacterial evolution. This technology will enable stakeholders to empirically evaluate the risks associated with resistance evolution before drug administration ***Rolff et al. (2024***). To reach this quixotic but worthwhile endeavor, we need to develop computational models and analytical tools to predict how antibiotic resistance will evolve ***Stracy et al. (2022***); ***Weinstein et al. (2001***).

Evolutionary therapies have been developed by applying optimization methods to mathematical models of the evolving system ***Goulart et al. (2013***); ***Hernandez-Vargas et al. (2013***); ***Nichol et al. (2015***); ***Mira et al. (2015***); ***Yoon et al. (2018***); ***Tetteh et al. (2020***); ***Nyhoegen and Uecker (2023***). Most antibiotic cycling strategies have been determined by linking dynamical models to fitness landscapes ***Toprak et al. (2012***); ***Goulart et al. (2013***); ***Visser and Krug (2014***); ***Nichol et al. (2015***); ***Mira et al. (2015***); ***Baym et al. (2016***); ***Gjini and Wood (2021***). Other promising approaches are those based on reinforced learning approaches, e.g., ***Weaver et al. (2024***), which are capable of learning effective drug cycling policies. However, most machine learning algorithms require large data sets, thus basing the training on dynamically measured fitness landscapes. Another key obstacle from previous computational approaches is the assumption of a singular protein landscape -the complexity of epistatic networks. While this can facilitate analysis by Markov models ***Goulart et al. (2013***) and other machine tools ***Weaver et al. (2024***), the evolutionary constraint is limited to the number of mutations.

Collateral sensitivity, a bacterium that has developed resistance to one antibiotic becomes more susceptible to another antibiotic, is a central biological mechanism to combat antibiotic resistance ***Pluchino et al. (2012***); ***Lázár et al. (2013***); ***Munck et al. (2014***); ***Imamovic and Sommer (2013***); ***Podnecky et al. (2018***). Cycling antibiotics based on collateral sensitivity presents an exceptional evolutionary therapy to tackle antibiotic resistance ***Imamovic and Sommer (2013***). However, scheduling the order and time of antibiotics by trial and error is not feasible.

Recent mathematical models ***Tetteh et al. (2023***); ***Aulin et al. (2021***); ***Nyhoegen and Uecker (2023***) show that cross-sensitivity-based dosing schedules could predict an effective antibiotic scheduling for suppression of within-host emergence of antibiotic resistance. The main drawback of previous modeling literature is that there are only hypothetical populations, *i*.*e*., no data-driven. In this paper, we conceive a mathematical abstraction and the respective free computational platform that performs data-driven prediction to highlight the failure of a combination of drugs, see **Figure 1**. To our knowledge, this is the first instance of a new way to navigate collateral sensitivity to streamline antibiotic cycling. While our platform has limitations in predicting success, drugs that would fail in this platform would definitely fail in a realistic scenario.

**Figure 1.**
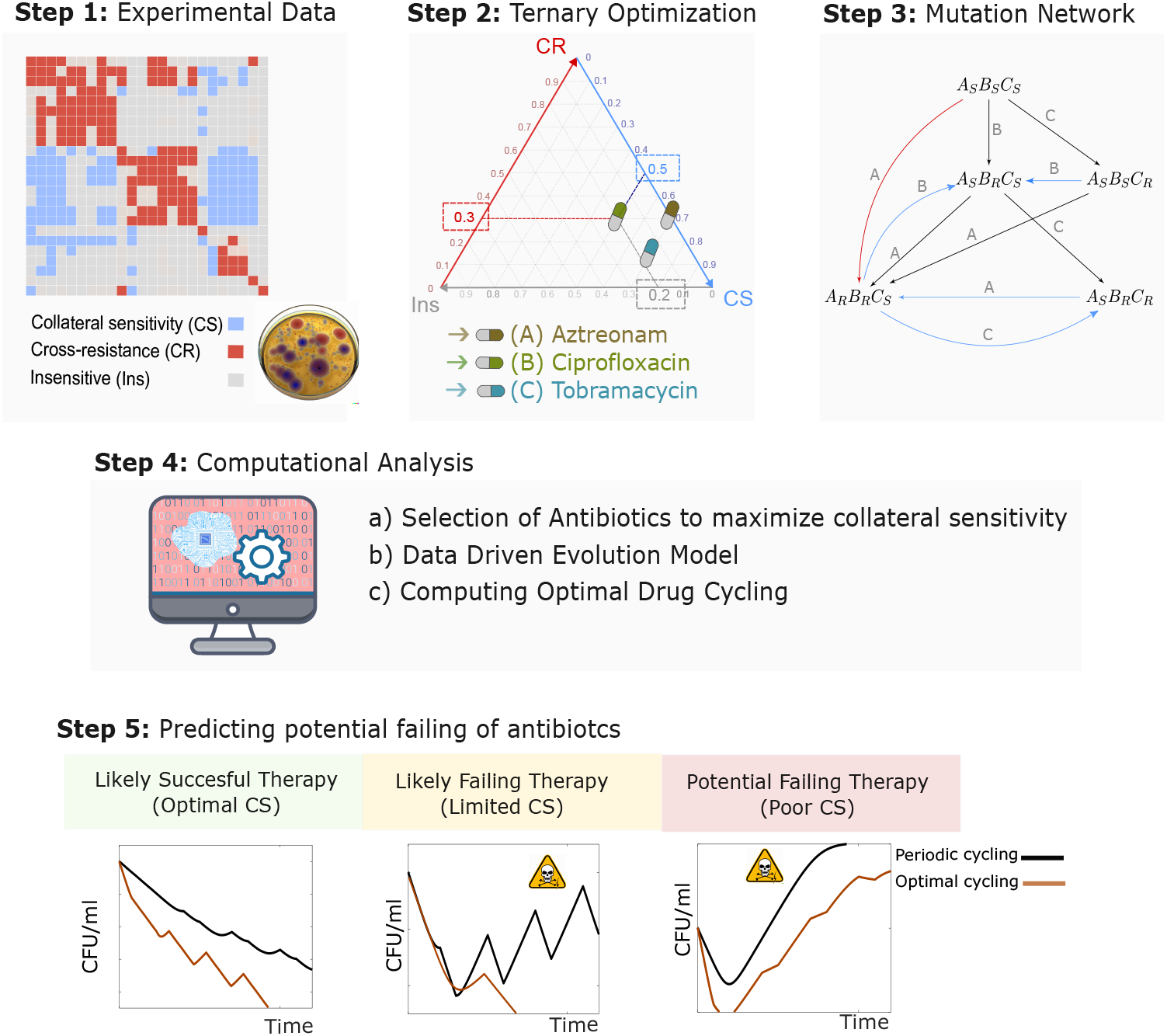
Intereractive Collateral Sensitivity Platform (ICSP). The computational platform requires the integration of quantitative MIC fold changes of the studied population as well as the selection of antibiotics to be evaluated (Panel A). Data with the profiles of a drug-evolved resistant strain is integrated into the computational platform (Panel B) to maximize collateral sensitivity in ternary plots (Panel B) and into a network model (Panel C). Ultimately, the network model is converted to a dynamical model to optimize cycling therapies.

## RESULTS

Our main result is the mathematical conception of collateral sensitivity integrated into an intuitive and easy used computational tool (**Figure 1**) that serves to *i)* avoid the selection of antibiotics that will trigger the emergence of a multi-resistance strain; *ii)* derive data-driven dynamical models to navigate a subspace of the evolution landscape; and *iii)* highlight antibiotic cycling failure.

It is worth emphasizing that our approach is based on a multivariable switched system of ordinary differential equations, which consider an instantaneous effect when a given drug is provided. While this platform is limited to providing the best sequence of drugs to eradicate a population, our predictions can indeed highlight which antibiotics and therapeutic sequences to avoid triggering resistance. This platform would offer a conservative scenario. That is, if a drug combination fails within our platform, the same combination would definitely fail if other more realistic conditions were included, such as pharmacodynamics and pharmacokinetics.

The applicability of our computational platform is presented next by integrating data of MIC fold changes of *Pseudomonas aeruginosa* (PA01) by ***Imamovic et al. (2018***) to 24 antibiotics (see Material and Methods section); red for MIC fold increase (cross-resistance, *CR*); blue for decrease (collateral sensitivity, *CS*); and gray for no change (insensitivity, *IN*).

### Modeling evolution effects of sequential antibiotics

By the concept of collateral sensitivity, we can assume that using antibiotic *A* can lead to the development of a strain that is susceptible to antibiotic *B*, despite being initially resistant to it. In this framework, any given collateral sensitivity, *CS*, causing suppressed resistance to another drug *R*, can be algebraically summarized as follows:

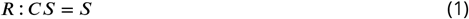

This is a nontrivial operation that requires careful consideration of the MIC FOLD differences needed to transition between what is defined as *R* (resistant) and *S* (sensitive) states. In Material and Methods we establish assumptions to fulfill six cross interactions *R* : *CS* = *S*; *R* : *CR* = *R*; *R* : *IN* = *R*; *S* : *CS* = *S*; *S* : *CR* = *R*; and *S* : *IN* = *S*, relative to states *R* and *S* for a variant. The evolutionary network in **Figure SC-B** was constructed by applying these operations to the wild-type strain (sensitive to all drugs) and subsequently to the emergent variants. An evolutionary network affecting all possible preexisting variants can be easily constructed as well, see **Figure SB** in Supplementary Material.

For example, **Figure SC** shows the effect of the interactions between antibiotics *Fosfomycin-FOS* (F), *Ceftazidime - CFZ* (C), *Amikacin-AMI* (A) and *Doxycycline-DOX* (D), on bacterial multiplication. The heatmap of these drugs can be observed in **Figure SC-A**. Observe how the wild-type strain *F*_*S*_*C*_*S*_*A*_*S*_*D*_*S*_ (sensitive to all) is affected by antibiotic AMI: because AMI shows *CR* towards DOX, the susceptibility of the wild-type to drug DOX, *F*_*S*_*C*_*S*_*A*_*S*_*D*_*S*_, develops resistance against DOX, *F*_?_*C*_?_*A*_*R*_*D*_*R*_, obtained by *S* : *CR* = *R*. Using the corresponding operation for drugs FOS and CFZ, a trend towards the strain *F*_*R*_*C*_*S*_*A*_*R*_*D*_*R*_ can be predicted, as depicted in **Figure SC-B**.

### Ternary diagrams for drug selection

Our computational platform provide ternary diagrams to give a straightforward visualization of antibiotic interactions, depicting proportions between collateral sensitivity, cross-resistance and insensitive. Our methodology effectively identifies drug combinations against the wild-type strain, but it can be extended to account for the presence of preexisting resistant strains (see Supplementary Material). **Figure 2** illustrates the solutions for three-drug therapeutic combinations across six different targets inside the ternary plot. We assess the total of 2, 024 possible combinations, among which 1, 485 resulted in treatment failure based on the escape criteria established earlier (see inset in **Figure 2**).

**Figure 2.**
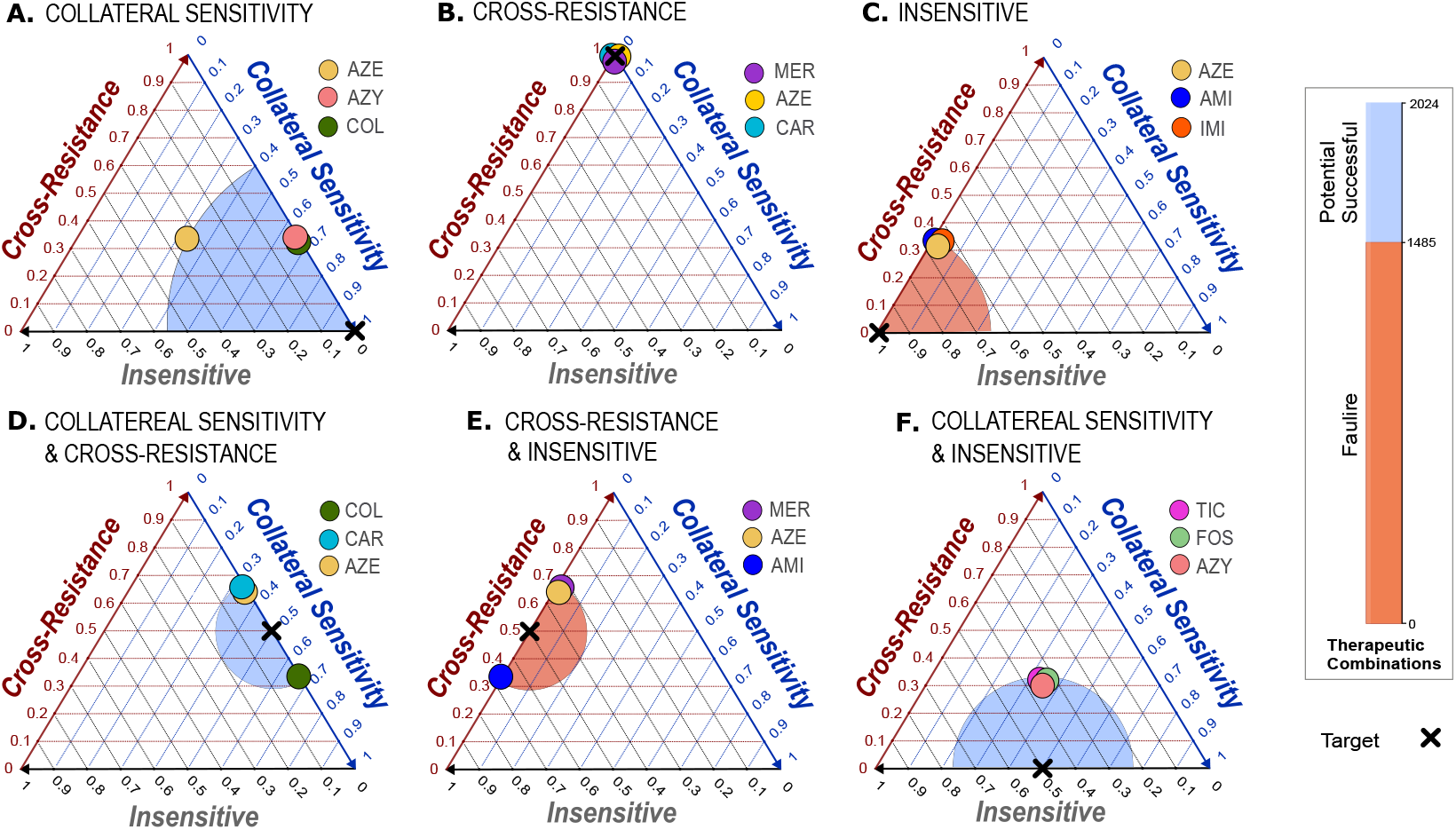
Ternary optimization for selection of three antibiotics. Three antibiotics entail the analysis of 2,024 different combinations. **A.** Antibiotics *Aztreonam* (AZE), *Azithromycin* (AZY) and *Colistin* (COL) is one solution exhibiting the highest level of collateral sensitivity, following a Target given by 100% of collateral sensitivity. **B**. *Meropenem* (MER), *Aztreonam* (AZE) and *Carbenicillin* (CAR) is among the solutions exhibiting the highest level of cross-resistance, following a Target of 100% cross-resistance. **C**. *Aztreonam* (AZE), *Amikacin* (AMI) and *Imipenem* (IMI) is one solution exhibiting the highest level of insensitive, with Target given by 100% insensitive. **D**. *Colistin* (COL), *Carbenicillin* (CAR) and *Aztreonam* (AZE) follows the Target given by 50% collateral sensitivity and 50% cross-resistance. **E**. *Meropenem* (MER), *Aztreonam* (AZE) and *Amikacin* (AMI) follows the Target given by 50% cross-resistance and 50% cross-resistance 50% insensitive. Finally, **F** *Ticarcillin* (TIC), *Fosfomycin* (FOS) and *Azithromycin* (AZY) follows the Target given by 50% collateral sensitivity and 50% insensitive. The inset displays the total of 2,024 combinations for a three drug therapy, out of which 1,485 are associated with therapeutic failure, leaving only 539 cases to analyze their success. The shaded area oximity represents the proximity of the solution to the target, red if the solution corresponds to therapy failure and blue otherwise.

The solutions in each panel on **Figure 2** converge towards the target as closely as possible, highlighted by the shaded area on each ternary plot (in red shade for the failed cases according to the established criteria and blue otherwise). **Figure 2-A** showcases a therapeutic combination closely aligning with the highest level of collateral sensitivity, marked by the Target 100%*CS*, 0%*CR*, and 0%*IN*. One solution includes *Aztreonam* (AZE), *Azithromycin* (AZY) and *Colistin* (COL). Notably, seven combinations out of the total 2, 024, exhibit identical interactions. The second panel, in **Figure 2-B**, identifies the set of antibiotics closer to the target 0%*CS*, 100%*CR*, and 0%*IN*. We show the solution *Meropenem* (MER), *Aztreonam* (AZE), and *Carbenicillin* (CAR), but there are 104 distinct combinations with the same interactions. The third panel, in **Figure 2-C**, shows that the set of antibiotics *Aztreonam* (AZE), *Amikacin* (AMI) and *Imipenem* (IMI), which demonstrates the higher level of insensitive. The Target here is 0%*CS*, 0%*CR* and 100%*IN*, and there are 26 combinations with the same interactions. In panel on **Figure 2-D**, the Target 50%*CS*, 50%*CR* and 0%*IN* yields 17 solutions with the same interaction as *Colistin* (COL), *Carbenicillin* (CAR) and *Aztreonam* (AZE). Panel in **Figure 2-E**, with Target 0%*CS*, 50%*CR* and 50%*IN* yields 133 solutions with the same interactions as *Meropenem* (MER), *Aztreonam* (AZE) and *Amikacin* (AMI). Finally, in **Figure 2-F**, the Target 50%*CS*, 0%*CR* and 50%*IN* yield 3 solutions with the interactions as *Ticarcillin* (TIC), *Fosfomycin* (FOS) and *Azithromycin* (AZY). As evident in the panels, the proximity of the solutions depends on the location of the target. Since all antibiotics display self-cross-resistance, there is a resistance trend, which can be observed in the proximity of the solutions. In Supplementary Material we show that increasing the number of antibiotics can mitigate this trend.

### Forecasting collateral effects in drug cycling

Our platform can integrate the evolutionary network into a dynamical system where the switch of drugs shifts the balance of growth and death such that the antibiotic-exposed population shrinks or escapes. This framework enables us to identify treatment failures and address the combinatorial complexity of drug combinations by effectively narrowing down a vast and counterintuitive array of ineffective potential combinations.

**Figure 3**. demonstrates how drug cycling affects bacterial growth, beginning with the wild-type strain PA01. Initially, the population shrinks regardless of the antibiotic used. However, after a few days, a rescue trajectory is noticeable in response to the stressors. More importantly, the response depends substantially on the interaction between antibiotics. **Figure 3-A** depicts a cycling regimen involving antibiotics on panel **Figure 2-A**: AZE, AZY and COL. The negative feedback in bacterial growth, given by a progressively declining of population during the cycling treatment, is associated with the synergy among these antibiotics. A minor rebound is observed between days 18 and 21, because AZE shows lower collateral sensitivity (according **Figure 2-A**). **Figure 3-B** shows a cycling regimen with antibiotics on panel **Figure 2-B**: MER, AZE and CAR. These exhibit the highest degree of cross-resistance, leading to rapid evolution of resistance, and consequently bacteria reaches the carrying capacity after 21 days of exposition. This failure was predicted by the established criterion on **Figure SD-A. Figure 3-C** shows a cycling regimen with antibiotics on panel **Figure 2-C**: AZE, AMI and IMI, which exhibit only insensitive interaction. There is a resistance delay, however this set of antibiotics lead to treatment failure (population shows positive feedback respect all antibiotics after day 18). This failure was predicted by the established criterion on **Figure SD-B. Figure 3-D** shows cycling regimen with antibiotics on panel **Figure 2-D**: COL, CAR and AZE, with cross-resistance and collateral sensitivity interactions respectively. Bacteria evolves resistance toward CAR and AZE first, because COL displayes higher collateral sensitivity. **Figure 3-E** depicts cycling regimen for antibiotics on panel **Figure 2-E**: MER, AZE and AMI. This scenario is also linked with failure based on the drugs positions. And **Figure 3-F** shows cycling regimen for antibiotics on panel **Figure 2-F**: TIC, FOS and AZY. These drugs exhibit 33% collateral sensitivity, 33% cross-resistance and 33% insensitive each. Bacteria evolves resistance toward TIC and FOS first, however since all drugs has identical interactions, it is expected that bacterial escapes after a longer period of time.

Interactions between drugs can be linked with either negative or positive feedback in bacterial multiplication. Additionally, insensitive proves to be an undesired interaction in sequential strategy, since bacteria decrease load at the onset of treatment but accumulated resistance triggers the escape of population after a long period of time.

**Figure 3.**
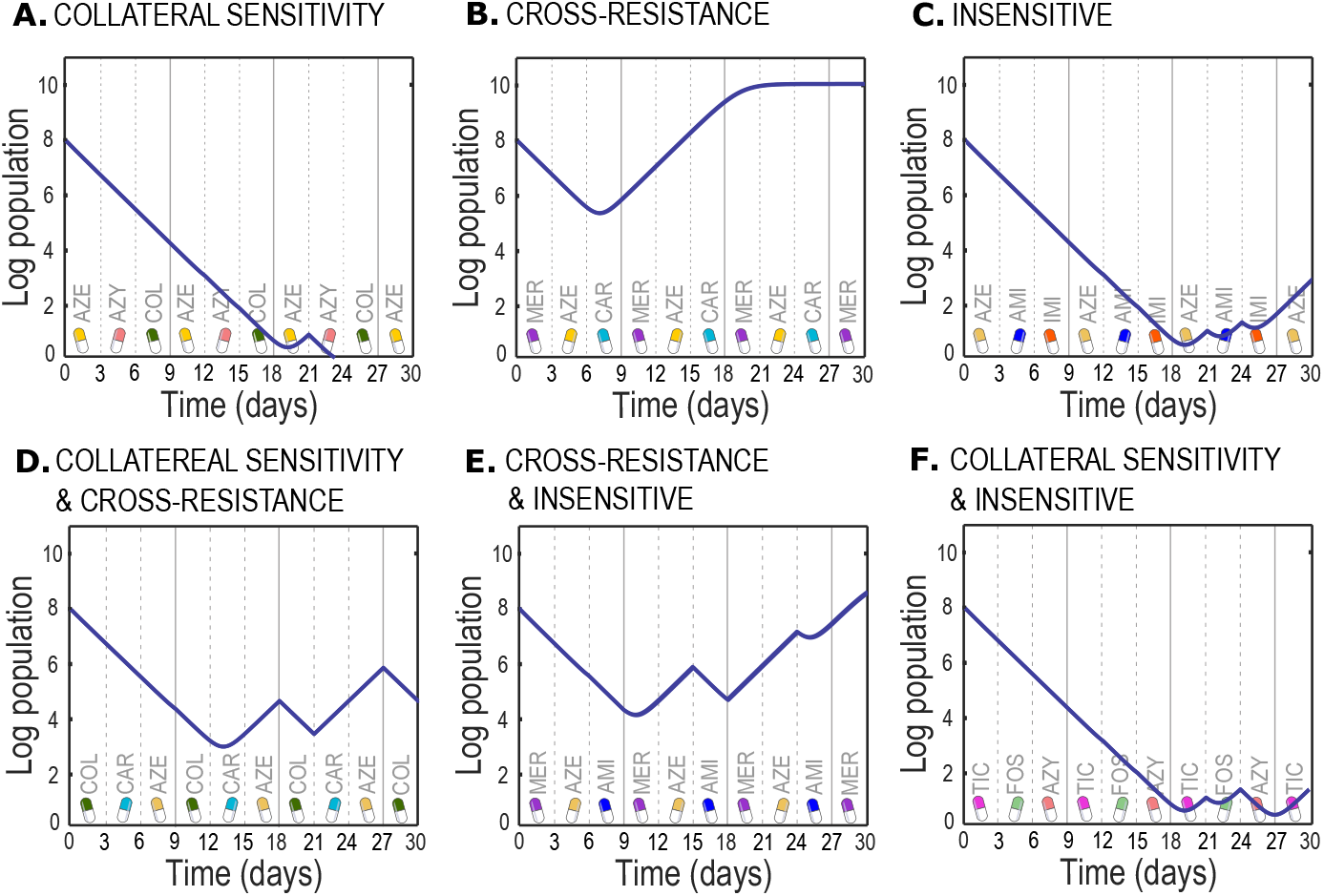
Qualitative long-term behavior of bacterial grow for cycling drugs. **A.** Evolution of PA01 under a cycling of antibiotics *Aztreonam* (AZE), *Azithromycin* (AZY) and *Colistin* (COL), which exhibit the highest level of collateral sensitivity. **B.** Prediction for cycling between *Meropenem* (MER), *Aztreonam* (AZE) and *Carbenicillin* (CAR), which exhibit the highest level of cross-resistance. **C.** Cycling antibiotics *Aztreonam* (AZE), *Amikacin* (AMI) and *Imipenem* (IMI) which show the higher level of insensitive interaction. **D.** Evolution of bacteria under cycling antibiotics *Colistin* (COL), *Carbenicillin* (CAR) and *Aztreonam* (AZE), which have 50% collateral sensitivity and 50% cross-resistance. **E.** Evolution of PA01 under a cycling of antibiotics *Meropenem* (MER), *Aztreonam* (AZE) and *Amikacin* (AMI), which have 50% cross-resistance and 50% insensitive. **F** shows bacterial growing for cycling *Ticarcillin* (TIC), *Fosfomycin* (FOS) and *Azithromycin* (AZY), which have have 50% collateral sensitivity and 50% insensitive.

### The impact of order in sequential antibiotic therapy

The careful selection of antibiotics to formulate treatment regimens is paramount, as it significantly influences therapeutic outcomes. However, simple decisions on how to apply these medications can also be decisive. **Figure 4** illustrate a dynamic control analysis designed to highlight the impact of order in drugs cycling. **Figure 4-D** shows that the inappropriate selection of drugs for a cycling, such as Ampicillin (AMI), Aztreonam (AZE) and Meropenem (MER), renders the order of antibiotics insignificant. Conversely, certain drug combinations exhibit enhanced efficacy when administered in a specific order, as demonstrated by simulations in **Figure 4-A**, for drugs Tobramycin (TOB), Ciprofloxacin (CIP) and Aztreonam (AZE). For a population, initially comprising by the wild-type (10^9^ number of bacteria) and each antibiotic exposure lasting 3 days per cycle for a total duration of 30 days, there is a pronounced difference in the susceptibility to a drug cycling designed to optimize killing rate, compared to a random drug cycling with the same antibiotics. The term *optimal* denotes a control strategy employed to determine the most effective order of antibiotic administration (refer to optimization details in Materials and Methods). These findings highlight that order is a significant factor in preventing the emergence of resistance and it can be decisive for the success or failure of a sequential strategies. In Supplementary Materials we explore other sub-optimal treatments by considering several factors.

**Figure 4.**
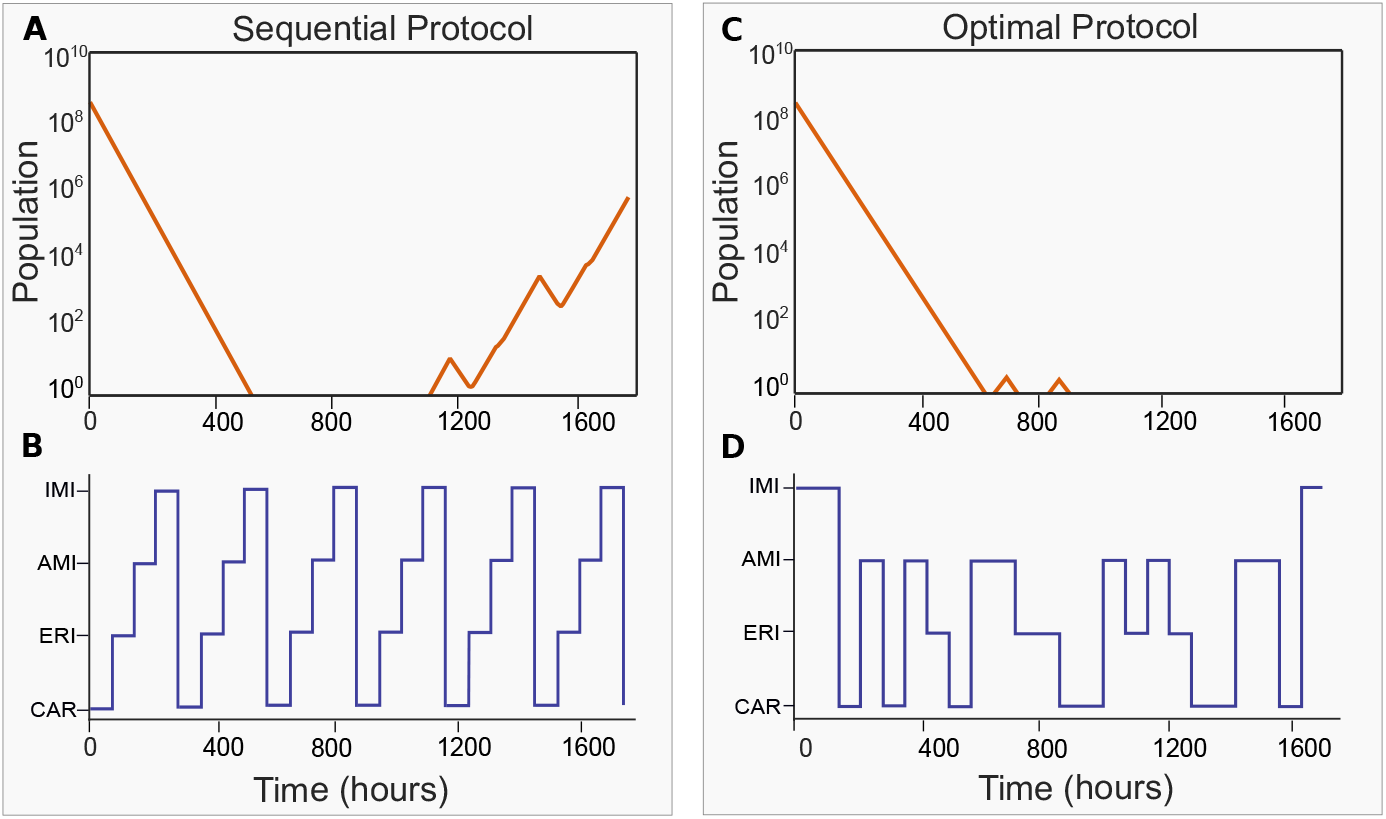
Total population dynamics for the sequential protocol (panel A and B) and optimal protocol (panel C and D).

## DISCUSSION

The discovery that a collateral sensitivity arises in bacterial populations following serial exposure to several antibiotics is a promising development in efforts to rationally design drug-cycling protocols. However, how best to apply the results of *in vitro* collateral sensitivity profiles obtaining for bacterial pathogens is debatable, and how laboratory experiments can be used to inform treatment protocols deserves broader analysis. This need can be attributed at least in part to difficulty in translating such data.

In this study we present an alternative approach to deducing optimal drug combinations that maximize the likelihood of treatment success based on collateral sensitivity. Several studies have been carried out on the emergence of collateral sensitivity and CS-interacting drugs were identified for *P*.*aeruginosa* ***Imamovic et al. (2018***), *E. coli* ***Imamovic and Sommer (2013***)***Jahn et al. (2017***) ***Podnecky et al. (2018***) and *E*.*faecalis* ***Maltas and Wood (2019***). The predominant method of analysis has been a precursory application of network theory to identify CS interacting drugs and identify patterns of collateral sensitivity. While some common themes obtained in the above-cited studies, e.g. CS of aminoglycoside resistant mutants towards other antibiotics, the results are far from uniform.

To this end, we introduced a nuanced approach to classifying drug cycling combinations that demonstrate the highest probability of success in a serially treated population of bacteria. In sequence, we i) streamlined the analysis of collateral sensitivity data generated *in vitro*, simplifying experimental evolutionary outcomes defined by the mutation network and correlated MIC changes, ii) formalized algebraically the defined phenotypic states (R or S) for *n* ≥ 2 antibiotics, iii) simulated the population dynamics for each genotype:phenotype component contributing to each cycling protocol (summarized in Figure 2), and iv) computed ternary diagrams that identified optimal combinations of interacting antibiotics (Figures 3 and 4). We validated this approach using the commensal sometimes pathogen *Bacteroides thetaiotaomicron* who we evolved to resistance against four clinically relevant antibiotics and tested for collateral sensitivity against each other. This data served as sufficient to predict an optimal treatment strategy which we successfully tested experimentally. With these results in mind, we sought to facilitate the deployment of our tool by other researchers and even clinicians. Thus, we provide an interactive tool that allows the rapid integration of experimental data into the prediction of optimal antibiotic cycling protocols. The utility of this approach is its ability to distill only the relevant information from a given experimental dataset.

The network approach commonly utilized in related studies merely describes the interaction between drug-pairs, only casually referring to cycling protocols. While the magnitude of an infecting pathogen’s MIC increase is informative, even small increments reduce the effective therapeutic window. They can result in treatment failure or from the microbial population’s perspective, evolutionary escape. Here, we chose to simplify the experimental ‘training’ data ***Imamovic et al. (2018***) focusing solely on the qualitative drug susceptibility classification of resistant vs sensitive. In accordance with the non-trivial algebraic operation set out in Equation 1, we described the mathematical relationships between the various singly and multiply resistant components of a sequentially exposed population of an erstwhile sensitive (wt) bacteria. *This is also valid for cases where resistance is pre-existent within the population*. Coupling these defined states into our birth and death-driven simulations and following the emergent properties of the system for longer durations of time, we obtained a higher resolution of single and multiple drug-resistant populations in competition with the ancestral population. For some identified CS drug pairs in ***Imamovic et al. (2018***), extended simulations using our modified framework led to treatment failure *in silico*. While we do not consider pharmacokinetics and extent of collateral sensitivity as ***Aulin et al. (2021***), we obtain similar results for two-drug combinations.

Notably, they have broad applications in various fields of science and were already applied to Hardy-Weinberg analyses of population genetics ***Finetti (1926***). Sequential applications of different drugs in such cases do not guarantee treatment success as subsequent evolution of an infecting pathogen to multi-resistance is often anticipated. We explore the utility of collateral sensitivity networks for the assessment and prediction of optimal combinations of drugs that maximize pathogen clearance. To complement efforts to predict drug courses that can minimize the probability of resistance arousal, we simplified the interaction between ‘bug’ and ‘drugs’ to a basic algebraic framework where resistance arousal was deterministic.

In summary, our open-source computational platform, grounded in the principles of collateral sensitivity, offers a powerful tool for anticipating treatment failures in managing multidrugresistant bacterial infections. By predicting therapeutic failures and facilitating the selection of effective drug regimens, our platform provides a valuable step toward reducing antibiotic resistance in persistent infections.

## MATERIAL AND METHODS

### Collateral sensitivity profiling data

Our platform requires data in the form of a heat map showing drug susceptibility profiles of antibiotic-resistant relative to wild-type bacteria. For an illustrative example, we consider that data in ***Imamovic et al. (2018***) who conducted a series of controlled experimental studies using bacterial populations subjected to antibiotic pressure, see **Figure 5**. The data in ***Imamovic et al. (2018***) was generated by evolving Pseudomonas aeruginosa under sequential drug exposure to characterize the evolved resistance profile. Phenotypic susceptibility assays and whole-genome sequencing are used to identify mutations associated with resistance development. Subsequently, ***Imamovic et al. (2018***) tested the susceptibility of the resistant strains to alternative antibiotics, identifying instances of collateral sensitivity—where resistance to one drug increases susceptibility to another. These results align with previous studies in cancer treatment, where collateral sensitivity-informed therapy has shown promising outcomes in limiting tumor evolution and improving therapeutic efficacy. This experimental foundation supports the use of computational models to predict optimal antibiotic cycling strategies, guiding the development of evolutionary-informed treatment approaches to combat antibiotic resistance.

**Figure 5.**
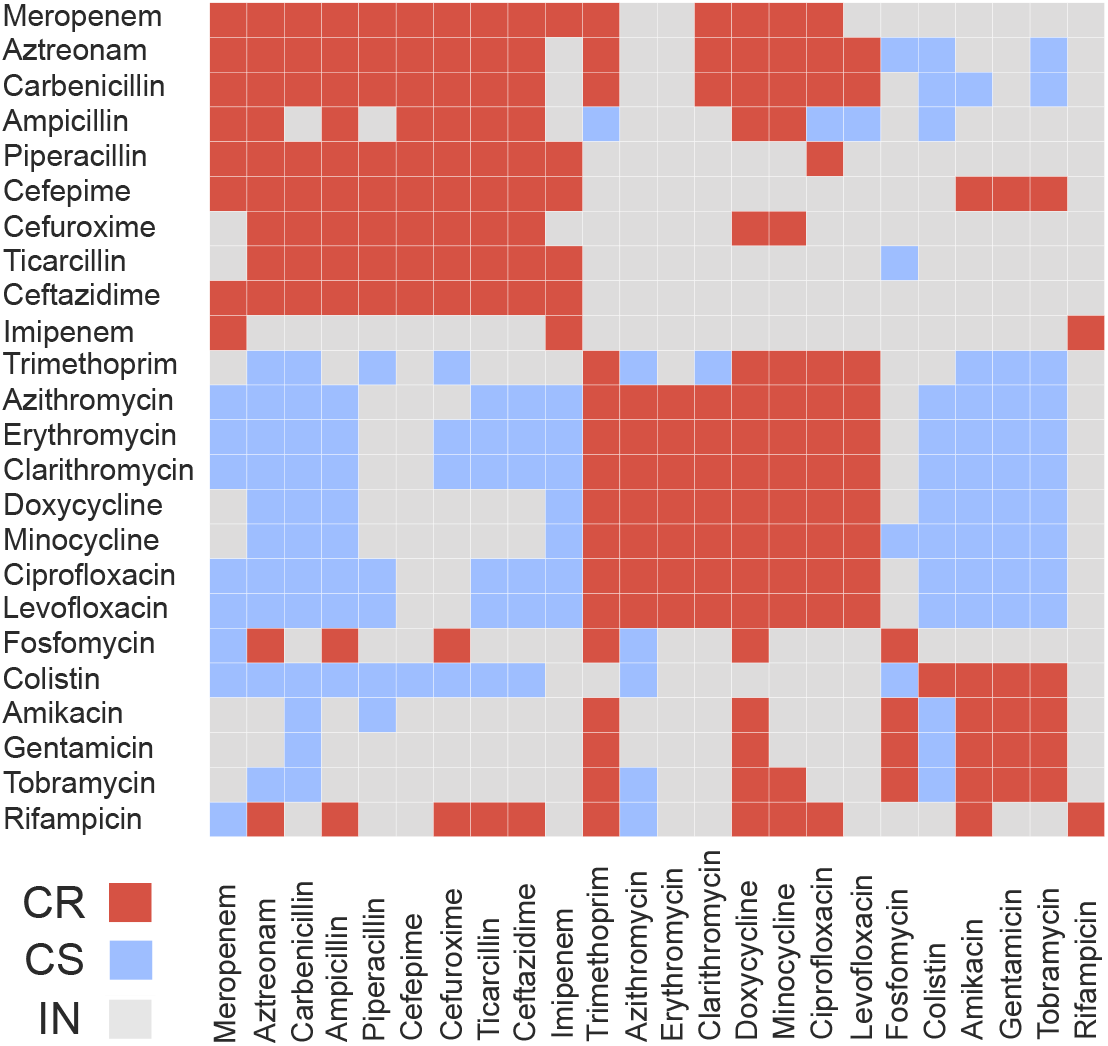
Antibiotic interactions for P. aeruginosa chronic infections: Qualitative data adapted from ***Imamovic et al. (2018***). A heatmap row delineates the profiles of a drug-evolved resistant strain. The interactions between the 24 antibiotics were examined by the effect on the wild-type PA01. Cross-resistance (CR) in red, collateral sensitivity (CS) in blue and insensitive (IN) in grey.

### Conceptualization of Sensitive (S) and Resistant (R) states for multiple antibiotics

Consider *k* antibiotics, {*σ*_1_, ⋯, *σ*_*k*_} = Σ, and a given concentration of a microorganism, *x*, the Minimum Inhibitory Concentration (MIC) parameter can be considered as *k*-dimensional as follows:

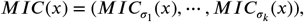

here, 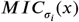 is the Minimum Inhibitory Concentration of drug *σ*_*i*_ for microorganism *x*. In addition, Breakpoints represent the maximum concentration of all drugs allowed for use, and can be represented by:

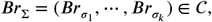

here, 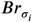 represents the maximum concentration of drug *σ*_*i*_ allowed for use, for all *i* = 1, ⋯, *k*, and 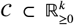 is the antibiotic concentration space. We can define the subsequent states for a microorganism *x* and antibiotics Σ.

#### Definition 1

(Sensitive/Resistant phenotypic state). *Consider a microorganism x and k antibiotics*, Σ = {*σ*_1_, ⋯, *σ*_*k*_}. *It is said that x is sensitive/resistant to drug σ*_*i*_, *with i* = 1, ⋯, *k, if the i*^*th*^ *element of the vector MIC*(*x*) − *Br*_Σ_, *is negative/non-negative*.

#### Definition 2

(Collateral effects). *Consider that state x*_*i*_ *converges phenotypically to state x*_*j*_ *under the pressure of drug* 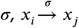. *Collateral effects can be measured by the vector:*

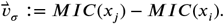

*If element i of* 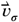 *is negative, then σ has* ***collateral sensitivity*** *against drug σ*_*i*_. *If element i is positive, then σ has* ***cross-resistance*** *against drug σ*_*i*_. *If element i of* 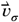 *is null, then σ has* ***insensitive*** *against drug σ*_*i*_.

For a set of *k* antibiotics, Σ = *σ*_1_, ⋯, *σ*_*k*_, and two susceptibility profiles (S and R) for microorganism *x* with respect to each antibiotic, there are 2^*k*^ possible states in the state-space model of Equation 4 in the main draft.

### Collateral sensitivity network

A model network structure is constructed by integrating the antibiotic interactions landscape of **Figure 6** as a flow diagram ***Blanchini and Giordano (2021***). The nodes of the network are binary strings of length equal to the number of drugs for cycling, let say *k* ≥ 2. The *σ*-th element of the node can be either sensitive (*S*) or resistant (*R*) to the *σ*-th drug of the list {1, 2, ⋯, *k*}. For *k* = 6 drugs, the node *RRRSSR* represents a microorganism resistant to drugs 1, 2, 3 and 6 but sensitive to drugs 4 and 5.

**Figure 6.**
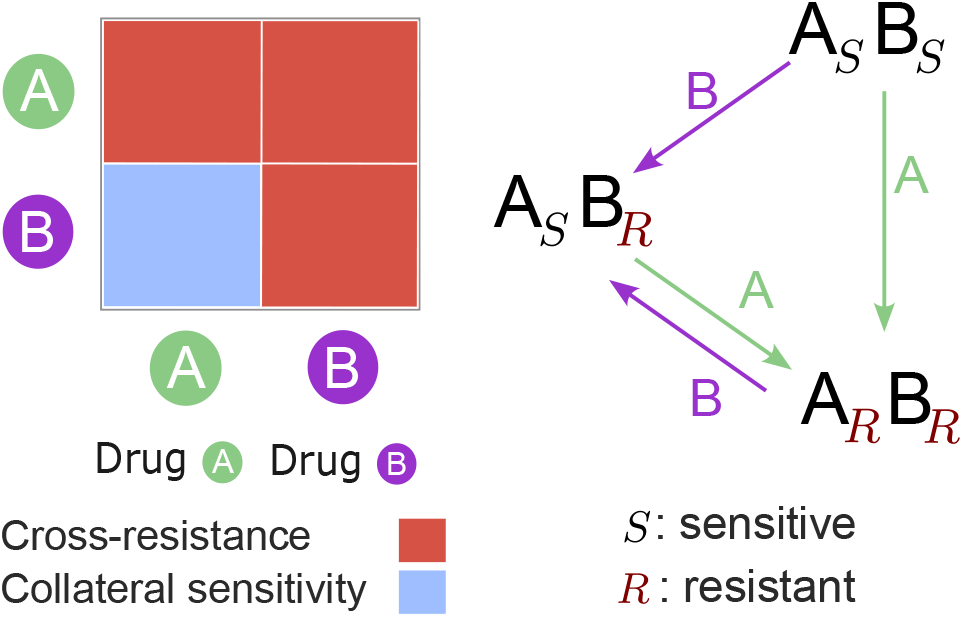
Drug *A* and drug *B* heatmap (left). Evolutionary network accounting for interaction between drugs *A* and *B* starting from the wild-type *A*_*S*_ *B*_*S*_ (right).

The interactions between antibiotics can be either collateral sensitivity (CS), cross-resistance (CR), or insensitivity (IN), and each can trigger a cross-proliferation rate between one node to another. The connections between nodes can be modeled through the following assumption:

#### Assumption 1.

*For bacteria resistant (R) to antibiotic B, which are stressed by antibiotic A, the following holds:*

*A.1.1. If A has CR respect to B, the bacteria mutate to a new variant resistant to B (R* : *CR* = *R)*.

*A.1.2. If A has CS respect to B, the bacteria mutate to a new variant sensitive to B (R* : *CS* = *S)*.

*A.1.3. If A has IN respect to B, the bacteria mutate to a new variant resistant to B (R* : *IN* = *R)*.

*For bacteria sensitive (S) to antibiotic B, which are stressed by antibiotic A, the following holds:*

*A.2.1.If A has CR respect to B, the bacteria mutate to a new variant resistant to B (S* : *CR* = *R)*.

*A.2.2. If A has CS respect to B, the bacteria mutate to a new variant sensitive to B (S* : *CS* = *S)*.

*A.2.3. If A has IN respect to B, the bacteria mutate to a new variant sensitive to B (S* : *IN* = *S)*.

There are 2^*k*^ nodes in a complete evolutionary network. However, in contrast to classical networks ***Komarova and Wodarz (2005***), not every node is necessarily present within our framework, some nodes may emerge from the preexisting variants, others not. The number of present nodes depends on the considered preexisting variants (or seeds of the network) and those emerging due to antibiotic interactions. **Figure 6** illustrates the evolutionary network based on the interactions between drug *A* and drug *B* and one seed, given by the wild-type *A*_*S*_*B*_*S*_ (sensitive to *A* and *B*). Applying drug *A* to the seed *A*_*S*_*B*_*S*_ directs the mutation towards node *A*_*R*_*B*_*R*_, in accordance with Assumption 1 (A.2.1). Applying drug *B* to the seed *A*_*S*_*B*_*S*_ directs the mutation towards node *A*_*S*_*B*_*R*_, as per Assumption 1 (A.2.2). Subsequently, applying drugs *A* and *B* to the emerging variants completes the mutational network. These operators provide a methodology to compute complete mutational network, for *k* > 1 drugs and with 2^*k*^ nodes, as presented in **Figure SB** in the Supplementary Material.

### Ternary diagram for selection of antibiotics

A ternary diagram is constructed to illustrates the proportion between Collateral Sensitivity (CS), Cross-Resistance (CR), and Insensitivity (IN) across a set of *k* antibiotics, **σ** = {*σ*_1_, ⋯, *σ*_*k*_}. Each drug *σ*_*i*_ will have *a*_*i*_ × 100% of CS, *b*_*i*_ × 100% of CR and *c*_*i*_ × 100% of IN, with *a*_*i*_ + *b*_*i*_ + *c*_*i*_ = 1. The position of antibiotic *σ*_*i*_ in the ternary diagram is given by 𝕋_*i*_ = (*a*_*i*_, *b*_*i*_, *c*_*i*_). **Figure 7** shows a heatmap for *k* = 5 hypothetical antibiotics and illustrates the construction of the ternary plot.

**Figure 7.**
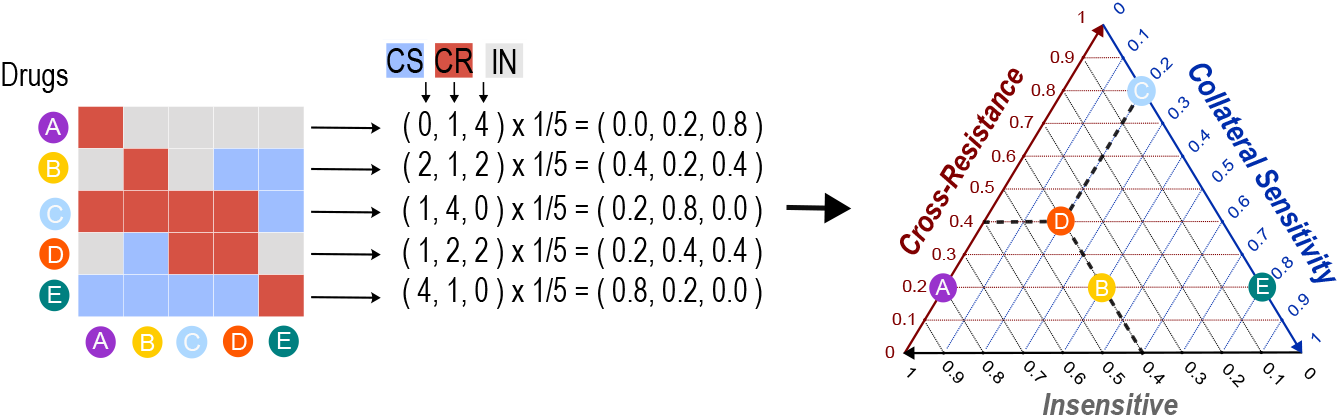
Ternary diagram construction.

We utilize the ternary diagram to design an antibiotic selection system. Assume we aim to select *k* antibiotics **σ** = {*σ*_1_, ⋯, *σ*_*k*_} from a list of *N* = 24 relevant antibiotics shown in **Figure 5**, such that the chosen *k* antibiotics are as proximal as possible to a specific target point within the ternary diagram, Target = (*T*_*cs*_, *T*_*cr*_, *T*_*in*_). The set *D*(*k, N*) represents the collection of all possible combinations of *k* antibiotics from the total *N* = 24, such that **σ** ∈ *D*(*k, N*) implies **σ** = {*σ*_1_, ⋯, *σ*_*k*_} conformed by different antibiotics from the set of *N* available antibiotics. Consequently, *D*(*k, N*) contains 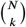 elements, where

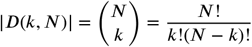

For any **σ** = {*σ*_1_, ⋯, *σ*_*k*_} ∈ *D*(*k, N*), we can compute the ternary position for every *σ*_*i*_ ∈ **σ**, represented by 𝕋_*i*_. Then, every sequence of *k* drugs, given by **σ**, has associated the following cost:

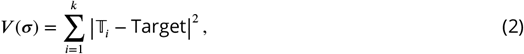

and we solve the optimization problem:

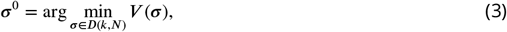

where **σ**^0^ is the sequence of *k* antibiotics closer to the Target = (*T*_*cs*_, *T*_*cr*_, *T*_*in*_) point inside the ternary diagram.

Selecting antibiotics entails evaluating 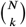 possible combinations and identifying the combination that minimizes the cost function *V* (*k*). The cost function (2) penalizes the sum of relative Euclidean distances between antibiotic positions against a target point within the ternary diagram. By employing this method, we can evaluate a vast number of combinations and optimize the set of antibiotics to match the specified level of interactions we need closely.

### Evolutionary dynamical model

The overall population dynamics are the outcome of two processes, the exponential decline of sensitive types (S) and the initially nearly exponential spread of resistant types (R). This process, called evolutionary rescue ***Bell (2017***); ***Ramsayer et al. (2013***); ***Drlica (2003***), delineates the foundational assumption upon which our dynamical analysis is built:

#### Assumption 2.

***(Evolutionary rescue)** In a sensitive bacterial population under antibiotic stress, the drug dose removes bacteria slowly enough for emerging resistant lineages to persist*.

The assumption above has been widely modeled using an ODE system ***Tetteh et al. (2023***). By considering two states, sensitive *S* and resistant *R*, with 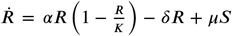, such that if there is an initial positive amount of state *S*, then state *R* emerges with a mutation rate *µ*. This will be used in the following section to model cycling antibiotics.

The concept of sequential drug combinations is integral to the switched system framework ***Liberzon (2003***); ***Hernandez-Vargas et al. (2011***), where transitions between drugs correspond to switch between systems. The nodes of the collateral sensitivity network, presented in the previous section, represent *n* different variants of a bacterium, which are the states of the switched system, given by *x*_*i*_, *i* = 1, …, *n*. Under pressure of antibiotic *σ* ∈ {1, 2, ⋯, *k*}, the rate at which the population *x*_*i*_ changes can be estimated by the addition of a growth, death and mutation terms. If 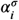 and 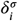 are growth and death rates of state *x*_*i*_, under pressure of drug *σ*, respectively, then we follows the next assumption:

#### Assumption 3.

*Variant x*_*i*_ *is sensitive to drug σ if and only if* 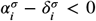, *and resistant if and only if* 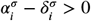.

For an algebraic formalization of defined sensitive and resistant phenotypic states for *k* ≥ 2 antibiotics see Definition 1 in Supplementary Material.

The collateral sensitivity network yields the matrix 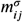 which represents the likelihood of a mutation from node *x*_*j*_ to node *x*_*i*_, under antibiotic *σ* exposure, with 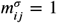 meaning the mutation is likely and 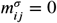 otherwise. The following equations estimate total bacterial size after a sequence of antibiotics given by the step function *σ*(*t*) : [0, ∞) → {1, 2, ⋯, *k*}:

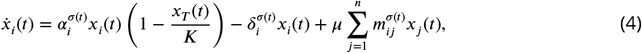

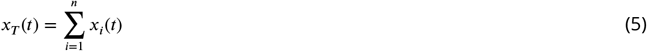

where 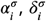, and *µ* are growth, death and mutation rates under pressure of drug *σ*, respectively. The state *x*_*T*_ denotes the total population, which growth strictly depends on the resistant strains as they serves as the exclusive mechanism for the growth of *x*_*T*_ . Saturation of *x*_*T*_ is given by saturated population density *K* (carrying capacity).

### Scheduling sequential antibiotics

From theory we know that a switched system, given solely by unstable subsystems, can stabilize the origin (see ***Anderson et al. (2021***) and references therein). This suggests that, despite Assumption 2, cyclic treatments have the potential to counteract the emergence of resistance.

To design a sequential therapy over the time interval [0, *T*_*f*_ ], *k* antibiotics must first be chosen. The switching times between one antibiotic and another are fixed and given by 0 = *T*_0_ < *T*_1_ < ⋯ < *T*_*f*−1_ < *T*_*f*_, such that one antibiotic *σ*_*i*_ ∈ {1, 2, ⋯, *k*} is applied during the time interval [*T*_*i*_, *T*_*i*+1_). Then, the switching law is given by *σ*(*t*) = *σ*_*i*_ for all *t* ∈ [*T*_*i*_, *T*_*i*+1_) for *i* = 0, 1, ⋯, *f* − 1. A switching law defined in this manner is said to be in the feasible set Σ. We want to find the switching law *σ*(*t*) ∈ Σ that minimizes the following cost function:

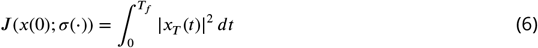

for a given initial condition *x*(0) = (*x*_1_(0), ⋯, *x*_*n*_(0)), and *x*_*T*_ (*t*) given by Eq. (5). To do this, we solve the following problem:

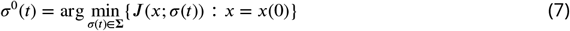

Note that *σ*^0^(*t*) is the optimal solution that minimizes the total population growth *x*_*T*_, considering only the order of the drugs. Other key factors in the control of switched systems include timing, dosage, consideration of limitations, and the order of drug sequences. The optimization is performed by using the DE algorithm ***Storn and Price (1997***), which is selected because of its simplicity and effectiveness.

### Software availability

We developed an open-source platform that provides an intuitive interface based on data on bacterial resistance profiles. Our tool enables rapid, data-driven decision-making to optimize therapeutic interventions. This can be found in:

https://github.com/matt0002/Collateral-Sensitivity-Networks.

## Acknowledgment of Support

This material is based upon work supported by the National Science Foundation under Grant No. 2315862.

## References

Anderson A, Gonzalez AH, Ferramosca A, Hernandez-Vargas EA. Discrete-time MPC for switched systems with applications to biomedical problems. Communications in Nonlinear Science and Numerical Simulation. 2021; 95:105586.

Annunziato G. Strategies to overcome antimicrobial resistance (AMR) making use of non-essential target inhibitors: A review. International journal of molecular sciences. 2019; 20(23):5844.

Aulin LB, Liakopoulos A, van der Graaf PH, Rozen DE, van Hasselt JC. Design principles of collateral sensitivity-based dosing strategies. Nature Communications. 2021; 12(1):5691.

Baym M, Stone LK, Kishony R. Multidrug evolutionary strategies to reverse antibiotic resistance. Science. 2016; 351(6268):aad3292.

Bell G. Evolutionary rescue. Annual Review of Ecology, Evolution, and Systematics. 2017; 48:605–627.

Blanchini F, Giordano G. Structural analysis in biology: A control-theoretic approach. Automatica. 2021; 126:109376.

Bollenbach T. Antimicrobial interactions: mechanisms and implications for drug discovery and resistance evolution. Current opinion in microbiology. 2015; 27:1–9.

Drlica K. The mutant selection window and antimicrobial resistance. Journal of Antimicrobial Chemotherapy. 2003; 52(1):11–17.

Finetti B. Considerazioni matematiche sull’ ereditarietà mendeliana. Metron. 1926; 6:29–37.

Gjini E, Wood KB. Price equation captures the role of drug interactions and collateral effects in the evolution of multidrug resistance. eLife. 2021 7; 10. doi: 10.7554/ELIFE.64851.

Gold HS, Moellering Jr RC. Antimicrobial-drug resistance. New England journal of medicine. 1996; 335(19):1445– 1453.

Goulart CP, Mahmudi M, Crona KA, Jacobs SD, Kallmann M, Hall BG, Greene DC, Barlow M. Designing antibiotic cycling strategies by determining and understanding local adaptive landscapes. PloS one. 2013; 8(2):e56040.

Hernandez-Vargas E, Colaneri P, Middleton R, Blanchini F. Discrete-time control for switched positive systems with application to mitigating viral escape. International journal of robust and nonlinear control. 2011; 21(10):1093–1111.

Hernandez-Vargas EA, Colaneri P, Middleton RH. Optimal therapy scheduling for a simplified HIV infection model. Automatica. 2013; 49:2874–2880.

Imamovic L, Ellabaan MMH, Machado AMD, Citterio L, Wulff T, Molin S, Johansen HK, Sommer MOA. Drug-driven phenotypic convergence supports rational treatment strategies of chronic infections. Cell. 2018; 172(1-2):121–134.

Imamovic L, Sommer MO. Use of collateral sensitivity networks to design drug cycling protocols that avoid resistance development. Science translational medicine. 2013; 5(204):204ra132–204ra132.

Jahn LJ, Munck C, Ellabaan MM, Sommer MO. Adaptive laboratory evolution of antibiotic resistance using different selection regimes lead to similar phenotypes and genotypes. Frontiers in Microbiology. 2017; 8(816):1– 14.

Komarova NL, Wodarz D. Drug resistance in cancer: principles of emergence and prevention. Proceedings of the National Academy of Sciences. 2005; 102(27):9714–9719.

Lázár V, Pal Singh G, Spohn R, Nagy I, Horváth B, Hrtyan M, Busa-Fekete R, Bogos B, Méhi O, Csörgő B, et al. Bacterial evolution of antibiotic hypersensitivity. Molecular systems biology. 2013; 9(1):700.

Levy SB. The challenge of antibiotic resistance. Scientific American. 1998; 278(3):46–53.

Levy SB, Marshall B. Antibacterial resistance worldwide: causes, challenges and responses. Nature medicine. 2004; 10(Suppl 12):S122–S129.

Liberzon D. Switching in systems and control. Springer Science & Business Media; 2003.

Maltas J, Wood KB. Pervasive and diverse collateral sensitivity profiles inform optimal strategies to limit antibiotic resistance. PLOS Biology. 2019; 17(10).

Mayers DL, Sobel JD, Ouellette M, Kaye KS, Marchaim D. Antimicrobial Drug Resistance: Clinical and Epidemiological Aspects, Volume 2, vol. 2. Springer; 2017.

Mira PM, Crona K, Greene D, Meza JC, Sturmfels B, Barlow M. Rational design of antibiotic treatment plans: a treatment strategy for managing evolution and reversing resistance. PloS one. 2015; 10(5):e0122283.

Munck C, Gumpert HK, Wallin AIN, Wang HH, Sommer MO. Prediction of resistance development against drug combinations by collateral responses to component drugs. Science translational medicine. 2014; 6(262):262ra156–262ra156.

Murray CJ, Ikuta KS, Sharara F, Swetschinski L, Aguilar GR, Gray A, Han C, Bisignano C, Rao P, Wool E, et al. Global burden of bacterial antimicrobial resistance in 2019: a systematic analysis. The lancet. 2022; 399(10325):629– 655.

Nichol D, Jeavons P, Fletcher AG, Bonomo RA, Maini PK, Paul JL, Gatenby RA, Anderson ARA, Scott JG. Steering Evolution with Sequential Therapy to Prevent the Emergence of Bacterial Antibiotic Resistance. PLOS Computational Biology. 2015; 11:e1004493. https://journals.plos.org/ploscompbiol/article?id=10.1371/journal.pcbi.1004493, doi: 10.1371/JOURNAL.PCBI.1004493.

Nyhoegen C, Uecker H. Sequential antibiotic therapy in the laboratory and in the patient. Journal of the Royal Society Interface. 2023 1; 20. https://royalsocietypublishing.org/doi/10.1098/rsif.2022.0793, doi: 10.1098/RSIF.2022.0793.

Perry GH. Evolutionary medicine. eLife. 2021 7; 10. doi: 10.7554/ELIFE.69398.

Pluchino KM, Hall MD, Goldsborough AS, Callaghan R, Gottesman MM. Collateral sensitivity as a strategy against cancer multidrug resistance. Drug Resistance Updates. 2012; 15(1-2):98–105.

Podnecky NL, Fredheim EGA, Kloos J, Sørum V, Primicerio R, Roberts AP, Rozen DE, Samuelsen Johnsen PJ. Conserved collateral antibiotic susceptibility networks in diverse clinical strains of Escherichia coli. Nature Communications 2018 9:1. 2018 9; 9:1–11. https://www.nature.com/articles/s41467-018-06143-y, doi: 10.1038/s41467-018-06143-y.

Ramsayer J, Kaltz O, Hochberg ME. Evolutionary rescue in populations of Pseudomonas fluorescens across an antibiotic gradient. Evolutionary applications. 2013; 6(4):608–616.

Rolff J, Bonhoeffer S, Kloft C, Leistner R, Regoes R, Hochberg ME. Forecasting antimicrobial resistance evolution. Trends in Microbiology. 2024; .

Storn R, Price K. Differential evolution–a simple and efficient heuristic for global optimization over continuous spaces. Journal of global optimization. 1997; p. 341–359. doi: 10.1023/A:1008202821328.

Stracy M, Snitser O, Yelin I, Amer Y, Parizade M, Katz R, Rimler G, Wolf T, Herzel E, Koren G, Kuint J, Foxman B, Chodick G, Shalev V, Kishony R. Minimizing treatment-induced emergence of antibiotic resistance in bacterial infections. Science. 2022 2; 375:889–894. https://www.science.org, doi: 10.1126/SCIENCE.ABG9868.

Tetteh JNA, Matthäus F, Hernandez-Vargas EA. A survey of within-host and between-hosts modelling for antibiotic resistance. Biosystems. 2020 10; 196:104182. doi: 10.1016/J.BIOSYSTEMS.2020.104182.

Tetteh JN, Olaru S, Crauel H, Hernandez-Vargas EA. Scheduling collateral sensitivity-based cycling therapies toward eradication of drug-resistant infections. International Journal of Robust and Nonlinear Control. 2023; 33(9):4824–4842.

Toprak E, Veres A, Michel JB, Chait R, Hartl DL, Kishony R. Evolutionary paths to antibiotic resistance under dynamically sustained drug stress. Nature genetics. 2012 1; 44:101. /pmc/articles/PMC3534735//pmc/articles/PMC3534735/?report=abstracthttps://www.ncbi.nlm.nih.gov/pmc/articles/PMC3534735/, doi: 10.1038/NG.1034.

Vestergaard M, Paulander W, Marvig RL, Clasen J, Jochumsen N, Molin S, Jelsbak L, Ingmer H, Folkesson A. Antibiotic combination therapy can select for broad-spectrum multidrug resistance in Pseudomonas aeruginosa. International journal of antimicrobial agents. 2016; 47(1):48–55.

Visser JAGMD, Krug J. Empirical fitness landscapes and the predictability of evolution. Nature Reviews Genetics 2014 15:7. 2014 6; 15:480–490. https://www.nature.com/articles/nrg3744, doi: 10.1038/nrg3744.

Weaver DT, King ES, Maltas J, Scott JG. Reinforcement learning informs optimal treatment strategies to limit antibiotic resistance. Proceedings of the National Academy of Sciences of the United States of America. 2024 4; 121. doi: 10.1073/PNAS.2303165121/SUPPL_FILE/PNAS.2303165121.SAPP.PDF.

Weinstein RA, Bonten MJ, Austin DJ, Lipsitch M. Understanding the spread of antibiotic resistant pathogens in hospitals: mathematical models as tools for control. Clinical Infectious Diseases. 2001; 33(10):1739–1746.

Yoon N, Velde RV, Marusyk A, Scott JG. Optimal Therapy Scheduling Based on a Pair of Collaterally Sensitive Drugs. Bulletin of mathematical biology. 2018 7; 80:1776–1809. https://pubmed.ncbi.nlm.nih.gov/29736596/, doi: 10.1007/S11538-018-0434-2.

