## Supplemental for "Computational Platform for streamlining the success of sequential antibiotic therapy"

[1]Department of Mathematics and Statistical Science, University of Idaho, Idaho, United States; [2]Department of Biological Sciences, Bioinformatics and Computational Biology, University of Idaho, Idaho, United States; [3]University of Littoral, Institute of Technological Development for the Chemical Industry and National Scientific and Technical Research Council, Santa Fe, Argentina; [4]Institute for Modeling Collaboration and Innovation, University of Idaho, Moscow, 83844–1103, Idaho, USA

### A. Results on emerging multi-resistant states

As previously mentioned, for a set of $k$ antibiotics, there are $2^k$ states. Some states can be considered preexisting in the initial population, others emerge alongside resistance, while some states never exist. Assumption 1 from Material and Methods in the main draft, gives some basic operations to predict connections between states, according the interaction data of antibiotics. Also, Assumption 2, from Material and Methods in the main draft, provides a condition for evolutionary rescue or resistance emergence. Both Assumptions can be used to prove the following results for the evolution of the wild-type variant, stressed by a cyclic sequence of antibiotics.

**Lemma SI.** *Subjecting the wild-type to a cycling drug regimen, with at least one antibiotic with cross-resistance across all antibiotics, leads to the emergence of a multidrug-resistant variant.*

**Lemma SII.** *Subjecting the wild-type to a cycling drug regimen, with at most one antibiotic with collateral sensitivity across others antibiotics, leads to the emergence of a multidrug-resistant variant.*

We are going to prove Lemma SII (Lemma SI is easier to prove). For this, suppose we have $k$ antibiotics, and we start with the wild-type variant $\underbrace{SSS\cdots S}_{k}$. We assume there is only one antibiotic with $CS$ in relation to the other drugs and the remaining drugs only have $IN$ (since $CR$ represents the worst-case scenario). When applying this drug in the cycle, we suppose, without loss of generality, that it was applied last, resulting in the state $SSS\cdots SR$ emerging. When applying Drug 1, we have $SSS\cdots R \xrightarrow{Drug1} RSS\cdots R$, only develops resistance respect to itself. Then, when applying Drug 2 to the second node, we have $RSS\cdots R \xrightarrow{Drug2} RRS\cdots R$, as this drug cannot counteract the resistance to Drug 1. If we continue with this reasoning, we can prove that after applying the rest

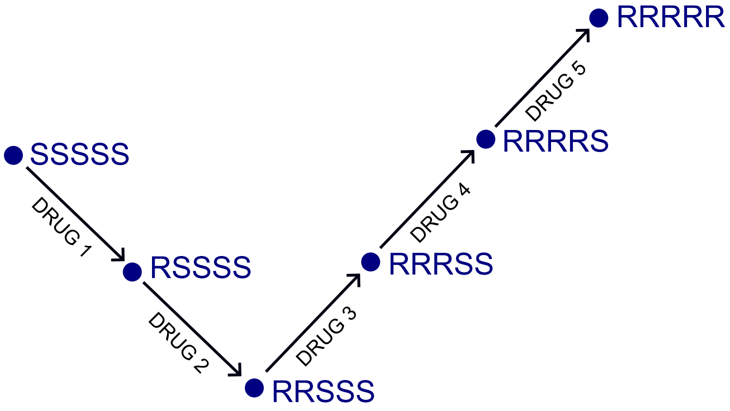


**Figure SA.** Cycling drugs facilitating an evolutionary pathway towards multi-resistance due to the absence of collateral sensitivities.

---

of the drugs to the corresponding node, the state $\underbrace{RRR\cdots R}_{k}$ emerges, which concludes the proof.

### B. Complete evolutionary networks

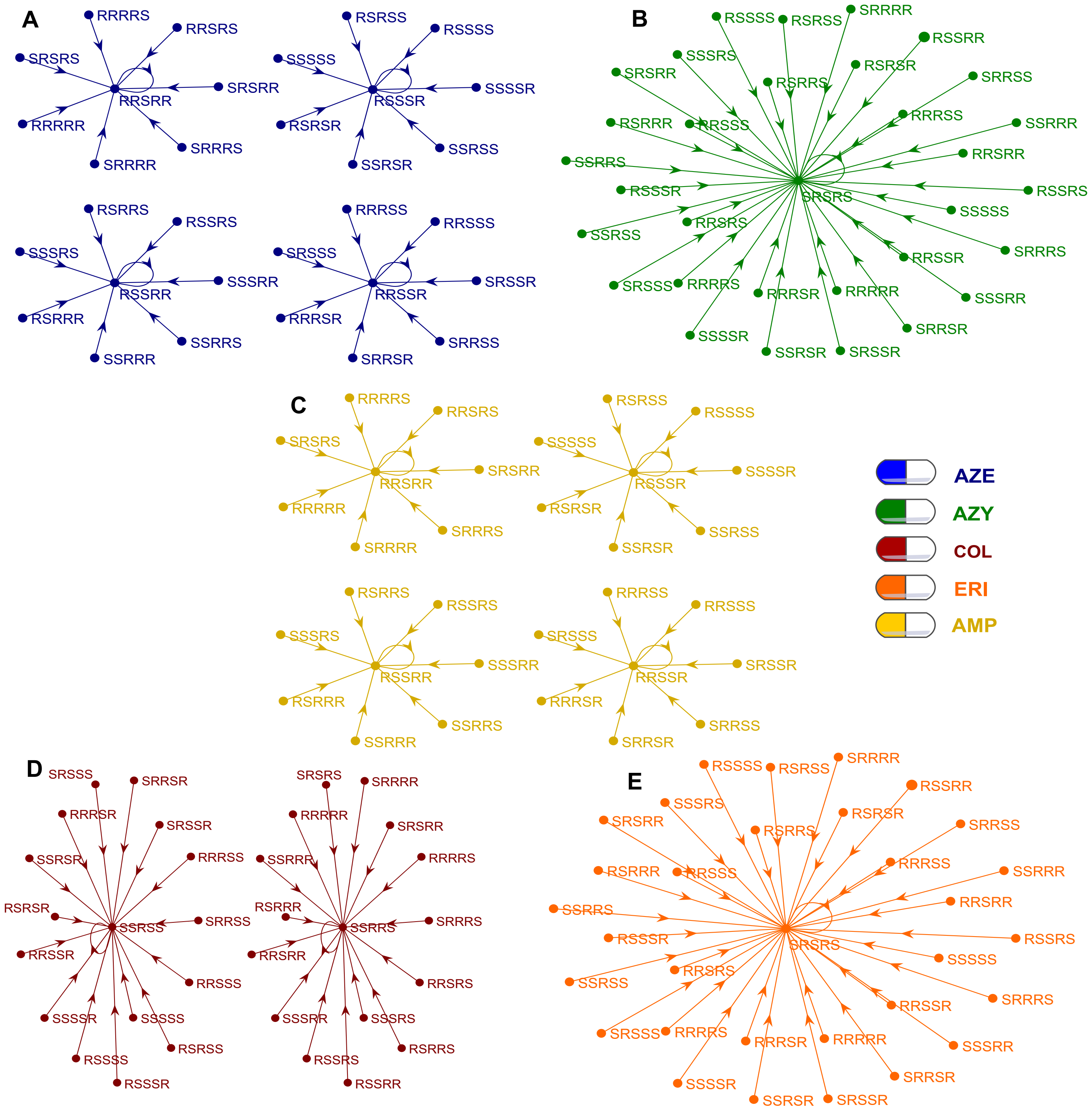


**Figure SB.** Complete evolutionary networks for P. aeruginosa variants according the interactions of antibiotics AZE (Aztreonam), AZY (Azithromycin), COL (Colistin), ERI (Erythromycin), AMP (Ampicillin) on Figure 5 in the main draft. For a set of five antibiotics, there are a total of $2^5 = 32$ possible states. We consider the evolutionary network between drugs $\Sigma = AZE, AZY, COL, ERI, AMP$, in such a way the variant $SSSRR$, represent a variant sensitive to the first three drugs in $\Sigma$, and resistant to the last two drugs in $\Sigma$. **A.**. Network in action under AZE drug pressure: describes the phenotypic evolution of each variant under the influence of the AZE drug. **B.**. Network under AZY drug pressure. **C.**. Network under COL drug pressure. **D.** Network under ERI drug pressure. **E.** Network under AMP drug pressure.

### C. Antibiotic Selection and Ternary plots

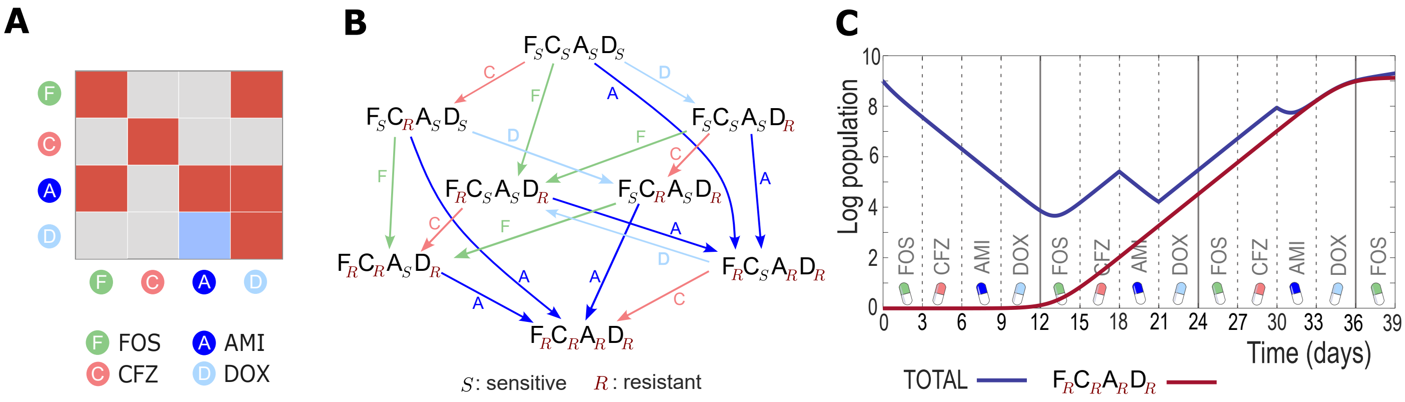


**Figure SC. Wrong antibiotics selection.** Cycling strategy for antibiotics *Fosfomycin* (FOS), *Ceftazidime* (CFZ), *Amikacin* (AMI), and *Doxycycline* (DOX). **A.** Heatmap of antibiotics $F = FOS$, $C = CFZ$, $A = AMI$ , and $D = DOX$, cross-resistance in red, collateral sensitivity in blue and insensitive in grey. **B.** Evolutionary network associated with interactions between drugs $F$, $C$, $A$ and $D$, for the wild-type $F_SC_SA_SD_S$ provided as the original population. The network uses collateral data to show how drug stress promotes the selection of a specific bacterial variant. **C.** Dynamic of total population $x_T$ (blue) and the multidrug resistant variant $F_RC_RA_RD_R$ (red), under cycling drugs. The cycle is defined by the ordered set $FOS - CFZ - AMI - DOX$, with a duration of 3 days per drug over a total exposure period of 39 days. The unavoidable escape of the population is attributed to variant $F_RC_RA_RD_R$. The multidrug-resistant variant $F_RC_RA_RD_R$ grows exponentially after 12 days of combination of antibiotics, leading to an unavoidable bacterial escape.

As an illustrative example, **Figure SC-C** depicts a computational output representing the resistance evolution of a population initially comprising by the wild-type ($10^9$ number of bacteria). Each antibiotic exposure lasting 3 days per cycle for a total duration of 39 days. Due to susceptibility, the total population, $x_T$, decreases until day 12. Subsequently, population $x_T$ develops resistance to the chosen cyclic treatment. Although still susceptible to drug AMI, as evidenced on day 18 when it is applied (this behavior is because the only collateral sensitivity present in this therapy is from antibiotic DOX respect to AMI). Exposition to DOX from day 21 onwards, causes an increase on population until switch to antibiotic AMI on day 30. At this point the population experiences a slight decline, but the accumulation of resistance in two nodes of the evolutionary network cause a rebounds and infection escapes. The simulations show that if AMI is placed at the end of the cycle, the effectiveness is improved, although the escape of population is unavoidable for a sequential therapy with this set of drugs. This set of drugs trigger the emergence of the multidrug resistant variant $F_RC_RA_RD_R$ according **Lemma SII**, this variant grows exponentially from day 12 onwards. We will later discuss how avoid these cases considering a ternary plot analysis, see this group of antibiotics represented in **Figure SD-B**.

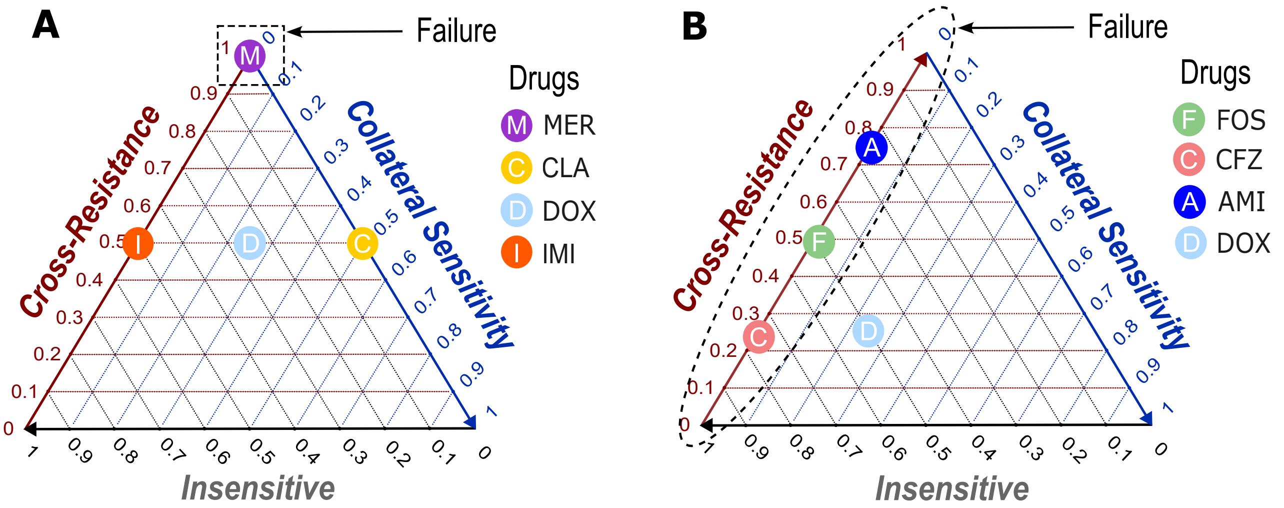


**Figure SD. Criteria for escape. A.** Ternary diagram for the drugs *Meropenem* (MER), *Clarithoromycin* (CLA), *Doxycycline* (DOX), and *Imipenem* (IMI). In this case, MER is in the region corresponding to $100\%$ of cross-resistance, which means it has $CR$ respect CLA, DOX, and IMI. Consequently, according to the dynamic model, any sequential therapy with these antibiotics leads to the escape of the total bacterial population. **B.** Ternary diagram for the drugs *Fosfomycin* (FOS), *Ceftazidime* (CFZ), *Amikacin* (AMI), and *Doxycycline* (DOX). In this case, all drugs, except DOX, are in the region corresponding to $0\%$ of collateral sensitivity. For a sequential therapy with $k$ drugs, if $k-1$ antibiotics belong to this region, the escape of the bacterial population is unavoidable.

Our dynamical analysis anticipates that a drug combination containing at least one antibiotic with cross-resistance across all drugs will lead to saturation of the total population density. This scenario applies to antibiotics such as *Meropenem* (MER), *Clarithoromycin* (CLA), *Doxycycline* (DOX) and *Imipenem* (IMI) in **Figure SD-A**. Note that MER shows $100\%$ of cross-resistance (against all drugs). Importantly, for the design of a four-drug sequential therapy from a set of $24$ antibiotics, there are $10,626$ possible combinations, out of which $1,237$ are predicted to fail due to this criterion, easily noticeable in a ternary plot as shown in **Figure SD-A**.

We anticipate that drug combinations containing at most one antibiotic with collateral sensitivity across others will also lead to saturation of the total population density. For example, the combination of antibiotics *Fosfomycin* (FOS), *Ceftazidime* (CFZ), *Amikacin* (AMI) and *Doxycycline* (DOX), as depicted in **Figure SD-B**. The computational dynamic solution of this example is also simulated in **Figure SC**, for a given cycling drug strategy. This criterion predicts the failure of $4,367$ combinations out of the total $10,626$, and this case is also easily noticeable in a ternary plot as shown in **Figure SD-B**.

The insights from **Figure SD-A** align with existing literature, providing a rationale for avoiding combinations of antibiotics within the same pharmacological class. Conversely, **Figure SD-B** uncovers intriguing combinations while also eliminating a substantial portion of possibilities. Notably, accounting for both failure criteria, a total of $4,769$ instances of four-drug combinations failing are identified among the $10,626$ possible combinations, all readily discernible through a ternary plot graph. In **Figure SE** we use these criteria to analyze the number of discarded combinations in general sequential therapies.

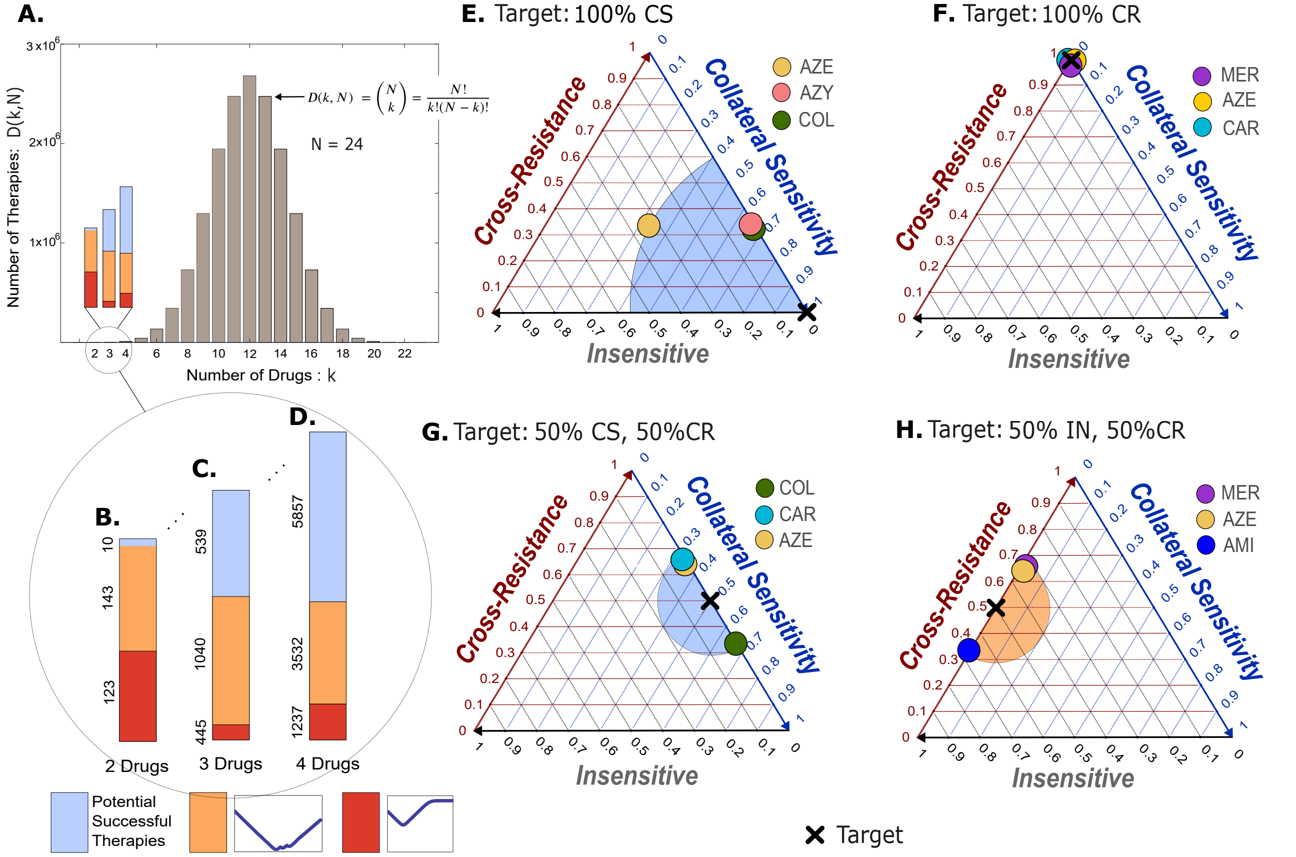


**Figure SE.** Selecting $k$ antibiotics from a list of $N$ antibiotics to optimize a cost (target within the ternary diagram) involves evaluating a large number of antibiotic combinations. For $N = 24$ drugs (the total number of antibiotics in the data shown in Figure 5), this number depends on $k$ and $N$ and is given by the formula $D(k, N) = \frac{N!}{k!(N-k)!}$. As shown in **A**, this grows rapidly with $k$ and has a maximum of $2,704,156$ for $k = 12$, then it decrease. **B.** Number of 2-drug combinations in the set of 24 drugs: 276. 123 combinations of these result in a failed therapy due to Lemma SI, 143 result in a failed therapy due to Lemma SII, and only 10 combinations have potential for success. **C.** Number of 3-drug combinations in the set of 24 drugs: 2024. 445 combinations of these result in a failed therapy due to Lemma SI, 1040 result in a failed therapy due to Lemma SII, and 539 combinations have potential for success. **D.** Number of 4-drug combinations in the set of 24 drugs: 10,626. 1237 combinations of these result in a failed therapy due to Lemma SI, 3532 result in a failed therapy due to Lemma SII, and 5857 combinations have potential for success. **E.** For the case of a 3-drug combination, we look for the one that most closely matches the target of $100\%\ CS$. One solution is given by the drugs $AZE$, $AZY$, and $COL$, with 7 total solutions exhibiting the same level of interactions, as shown in Table SA. **F.** For a 3-drug combination, *Meropenem* (MER), *Aztreonam* (AZE), and *Carbenicillin* (CAR) is among the solutions exhibiting the highest level of cross-resistance, following a target of $100\%$ cross-resistance. **G.** For a 3-drug combination, *Colistin* (COL), *Carbenicillin* (CAR), and *Aztreonam* (AZE) follow the target given by $50\%$ collateral sensitivity and $50\%$ cross-resistance. **H.** For a 3-drug combination, *Meropenem* (MER), *Aztreonam* (AZE), and *Amikacin* (AMI) follow the target given by $50\%$ cross-resistance and $50\%$ insensitivity.

**Table SA.** All combinations of three antibiotics with a maximum of mutual $CS$ associated with dynamics in Figure 3-A

| | | | |
|---|---|---|---|
| $AZE, AZY, COL$ | $AZE, AZY, TOB$ | $CAR, AZY, COL$ | $CAR, AZY, TOB$ |
| $AMP, AZY, COL$ | $AMP, CIP, COL$ | $AMP, LEV, COL$ | – |

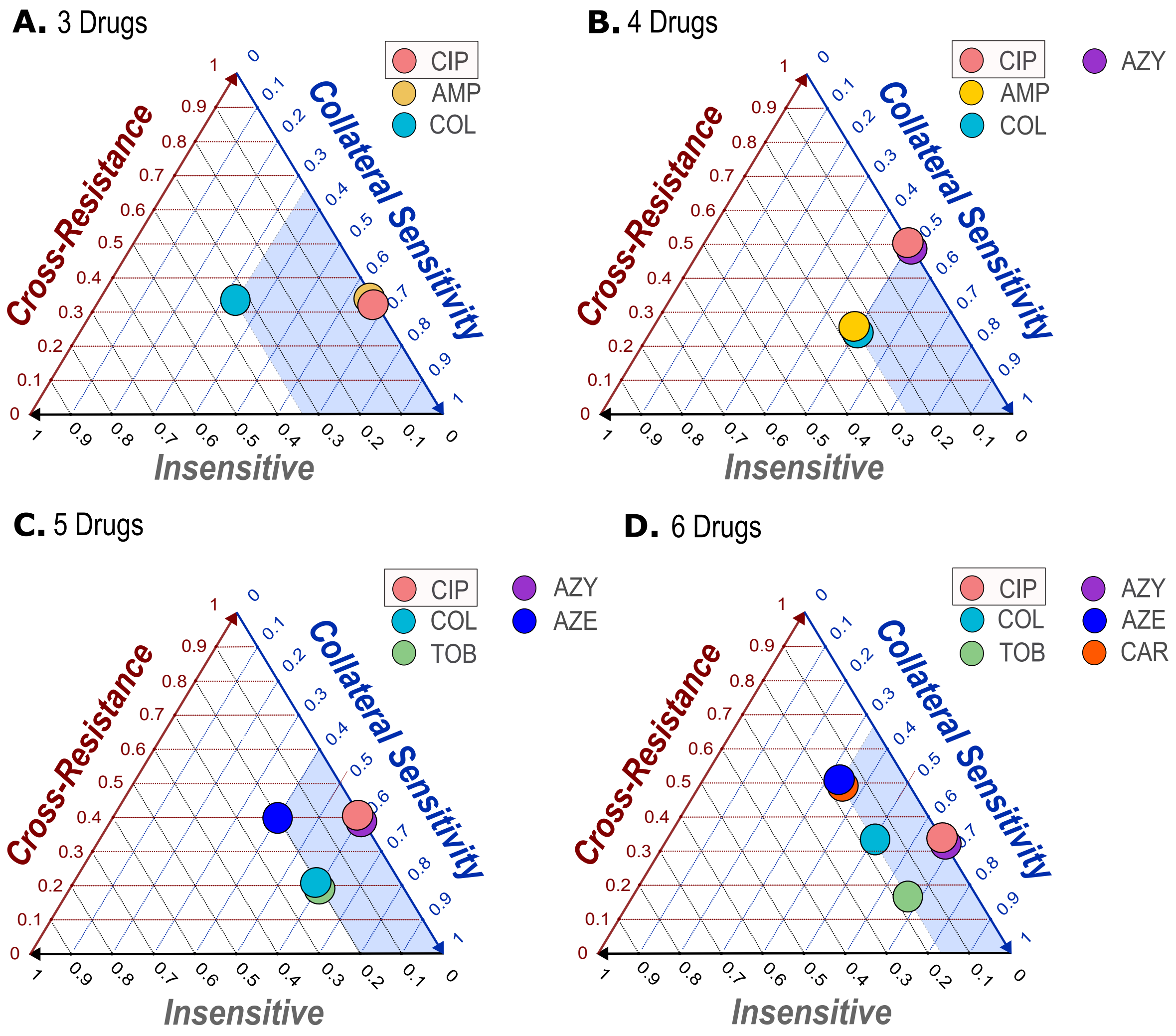


**Figure SF. The best combination of antibiotics for Ciprofloxacin (CIP)** was determined using ternary optimization. We searched for sets of 3, 4, 5, and 6 drugs that synergize best with Ciprofloxacin, aiming to maximize the percentage of collateral sensitivity of the set. The optimization utilized the target specified by $(CS, CR, IN) = (1, 0, 0)$
